# The extracellular matrix differentially directs myoblast motility in distinct forms of muscular dystrophy

**DOI:** 10.1101/2022.10.31.514525

**Authors:** Ashlee M. Long, Jason M. Kwon, GaHyun Lee, Nina. L. Reiser, Lauren. A Vaught, Joseph G. O’Brien, Patrick G.T. Page, Michele Hadhazy, Joseph C. Reynolds, Rachelle H. Crosbie, Alexis R. Demonbreun, Elizabeth M. McNally

**Affiliations:** Center for Genetic Medicine, Northwestern University Feinberg School of Medicine, Chicago, IL 60611, USA; Department of Integrative Biology and Physiology, UCLA, Los Angeles, CA; Department of Neurology David Geffen School of Medicine, UCLA, Los Angeles, CA; Department of Pharmacology, Northwestern University Feinberg School of Medicine, Chicago, IL 60611, USA

**Author notes:** To whom correspondence should be addressed: Alexis R. Demonbreun PhD Center for Genetic Medicine Northwestern University 303 E Superior SQ 5-512 Chicago, Il 60611 USA. Elizabeth M. McNally MD PhD Center for Genetic Medicine Northwestern University 303 E Superior SQ 5-516 Chicago, Il 60611 USA.

**Keywords:** Extracellular matrix, Duchenne muscular dystrophy, Limb Girdle muscular dystrophy, muscle, annexin, myoblast, Dysferlin

## Abstract

Extracellular matrix (ECM) pathologic remodeling underlies many fibrotic disorders, including muscular dystrophy. Tissue decellularization removes cellular components while leaving behind ECM components. We generated “on-slide” decellularized tissue slices from genetically distinct dystrophic mouse models. The ECM of dystrophin-and sarcoglycan-deficient muscles had marked thrombospondin 4 deposition, while dysferlin-deficient muscle had excess decorin. Annexins A2 and A6 were present on all dystrophic decellularized ECMs, but annexin matrix deposition was excessive in dysferlin-deficient muscular dystrophy. Adeno-associated viral expression of annexin A6 specifically in muscle resulted in annexin A6 deposition throughout the ECM, indicating muscle as a source of this ECM protein. C2C12 myoblasts seeded onto decellularized matrices displayed differential myoblast mobility. Dystrophin-deficient decellularized matrices inhibited myoblast mobility while dysferlin-deficient decellularized matrices enhanced myoblast movement. Myoblasts treated with recombinant annexin A6 increased mobillity similar to that seen on dysferlin-deficient decellularized matrix. These findings demonstrate specific fibrotic signatures elicit effects on myoblast activity.

**TEASER:** Fibrosis in muscular dystrophy has differential effects on myoblasts

**HIGHLIGHTS:**

- Spatial architecture and composition of the ECM differ across genetically distinct forms of muscular dystrophy, especially with respect to Annexin A6 protein deposition
- Matrix from dystrophin-mediated muscular dystrophy inhibits myoblast movement
- Matrix from dysferlin-deficient muscular dystrophy promotes myoblast motility
- Annexin A6 was sufficient to enhance myoblast motility

## INTRODUCTION

Muscular dystrophies are defined by continual replacement of muscle fibers by fibrosis. Patterns of fibrosis appear similar across genetically distinct forms of muscular dystrophy. Mutations that disrupt the dystrophin glycoprotein complex (DGC) produce a fragile muscle plasma membrane. Duchenne muscular dystrophy (DMD) is caused by loss-of-function mutations in the dystrophin gene (Duan et al., 2021; Rando, 2001). Within the DGC is the sarcoglycan complex, and recessive loss-of-function mutations in genes encoding α-, β-, δ-, or γ-sarcoglycan result in Limb Girdle muscular dystrophy (LGMD) that resembles DMD (Liu et al., 2019; Nigro and Savarese, 2014; Taghizadeh et al., 2019). Excessive muscle contraction and trauma can also induce membrane rupture, requiring a quick and efficient means of repair to prevent cellular necrosis. Dysferlin is a calcium-dependent, phospholipid-binding protein that facilities muscle membrane repair (Bansal et al., 2003; Bashir et al., 1998; Davis et al., 2002; Defour et al., 2014; Lek et al., 2012; McDade et al., 2014), and loss-of-function mutations in *DYSF*, the gene encoding dysferlin, also cause LGMD (Bashir *et al*., 1998; Liu et al., 1998). Dysferlin mutations are associated with remarkably high serum creatine kinase levels, indicative of muscle membrane leak, as well as proximal muscle weakness and inflammatory infiltrate (Contreras-Cubas et al., 2022). Patients with *DYSF* mutations, unlike other forms of muscular dystrophy, can often have completely normal to enhanced muscle function in early life with later onset of muscle weakness (Klinge et al., 2010).

Mouse models recapitulate many of the pathological features seen in the human muscular dystrophies (Ceco et al., 2014; Coley et al., 2016; Hack et al., 1998; Hammers et al., 2020; Ho et al., 2004; Lostal et al., 2010; Sicinski et al., 1989; van Putten et al., 2020). The *mdx* mouse harbors a single point mutation in exon 23, resulting in loss of full-length dystrophin (Sicinski *et al*., 1989). Genetic background can shift the pathological and functional manifestations of muscular dystrophy (Gordish-Dressman et al., 2018; Heydemann et al., 2005). Most mouse models, including *mdx* mice, are on C57BL substrains, and in this background these models exhibit relatively mild muscle disease. In contrast, the DBA/2J strain intensifies membrane fragility and fibrosis in *Sgcg null* mice, which lack the dystrophin-associated protein g-sarcoglycan (Heydemann *et al*., 2005). Similarly, in the DBA/2J strain the *mdx* mouse displays greater muscle damage, increased inflammation and increased intramuscular fibrosis compared to the C57BL/10 strain (Aartsma-Rus and van Putten, 2014; Coley *et al*., 2016).

The increase in disease severity conferred by the DBA/2J background has been attributed to genetic modifiers including latent TGF-β binding protein 4 (LTBP4); LTBP4 and the modifier osteopontin (OPN/SPP1) converge on the TGF-β pathway in a feed forward cycle (Quattrocelli et al., 2017). *Anxa6*, encoding annexin A6, was also mapped as a genetic modifier of muscular dystrophy (Swaggart et al., 2014). Annexin A6 (ANXA6) is a known membrane repair protein that acts as a molecular band-aid, aiding in membrane resealing, improving repair capacity upon membrane injury (Boye et al., 2017; Croissant et al., 2020; Demonbreun et al., 2019a; Demonbreun et al., 2016; Swaggart *et al*., 2014). Within the muscle extracellular matrix (ECM), the role of ANXA6 is less defined but, as a protein class, annexins have been implicated in cellular migration and differentiation (Belvedere et al., 2014; Bizzarro et al., 2012; Garcia-Melero et al., 2016; Grewal et al., 2017).

Like other tissues, the muscle ECM influences myofiber function as well as muscle repair and regeneration. In healthy muscle, the ECM includes collagens, glycoproteins, and proteoglycans, and supporting signaling proteins like TGF-β. After muscle injury, ECM remodeling is part of repair and recovery process. In muscular dystrophy however, there is ongoing and repetitive injury concomitant with attempted repair (Dowling et al., 2021; Mazala et al., 2020; Rayagiri et al., 2018). The excess ECM formation in muscular dystrophy impedes myoblast differentiation and promotes fibroblast activity (Loreti and Sacco, 2022; Thomas et al., 2015). A number of studies have identified impaired or altered myoblast properties in muscular dystrophy (Blau et al., 1983; Demonbreun et al., 2011; Gosselin et al., 2022). Recently, a method for the generation of acellular myoscaffolds was described, and this approach relies on removing the cellular content from dystrophic muscle in an “on-slide” format (Stearns-Reider et al., 2023). Using these acellular myoscaffolds, laminin remodeling was identified as critical for the adhesion and differentiation of skeletal muscle precursor cells (Stearns-Reider *et al*., 2023).

We now evaluated the ECM composition across different mouse models of muscular dystrophy by generating decellularized ECMs (dECMs). We identified distinct patterns of ECM proteins, including both core matrisome and matrisome-associated proteins when comparing DGC-related dystrophy to dysferlin-related dystrophy. We found that ECM from the DBA/2J (D2) dystrophic background had the most significant increase in matrisomal protein expression of decorin, periostin, thrombospondin 4 and matrix metalloproteinase 9. Interestingly, dysferlin-null muscle displayed the highest expression of matrix-associated annexin A2 and annexin A6. AAV9 expression of annexin A6 specifically in muscle produced extracellular deposition of annexin A6 indicating that ECM-associated ANXA6 is derived, at least in part, from muscle cells. dECMs supported unique effects on myoblast motility. C2C12 myoblasts seeded onto dECMs from the *mdx* DBA/2J background inhibited myoblast movement and migration. Conversely, migration and movement were enhanced on the dysferlin-null dECM compared to wildtype and *mdx* dystrophic scaffolds. Exposing myoblasts to recombinant annexin A6 was sufficient to enhance myoblast migration and movement similar to dysferlin-null scaffolds. These data illustrate that differential protein deposition into the ECM mediates cellular crosstalk in the muscular dystrophies.

## RESULTS

### Generation of acellular myoscaffolds using detergent decellularization

Decellularized scaffolds from heart and muscle have been studied for their ability to support cellular regeneration, but most commonly these methods have been applied to whole organs or tissues (reviewed in (Tan et al., 2022)). Recently Stearns-Reider et al published an “on-slide” decellularization method using 30μm tissue sections exposed to 1% SDS for 40 mins to generate dECM myoscaffolds (Stearns-Reider *et al*., 2023). We adapted and further optimized the generation of dECM scaffolds to ensure reproducible removal of intracellular components and DNA while retaining ECM architecture and function. We evaluated a range of sodium dodecyl sulfate (SDS) exposure by incubating 25μM sections from WT and *mdx* quadriceps muscles in 1% SDS for 10-30 minutes. Removal of the cellular components after decellularization was assessed through hematoxylin and eosin (H&E) staining, with complete decellularization observed by 10 minutes (**Figure 1A**). Decellularization was attempted with less SDS (0.1%) for 0-60 minutes (**Figure 1B**). However, at the lower concentration, decellularization remained incomplete after 20-40 minutes. Complete decellularization was seen after a 60-minute incubation with 0.1% SDS (**Figure 1B**), but for efficiency in tissue processing the higher SDS concentration was used at the shorter incubation time.

**Figure 1.**
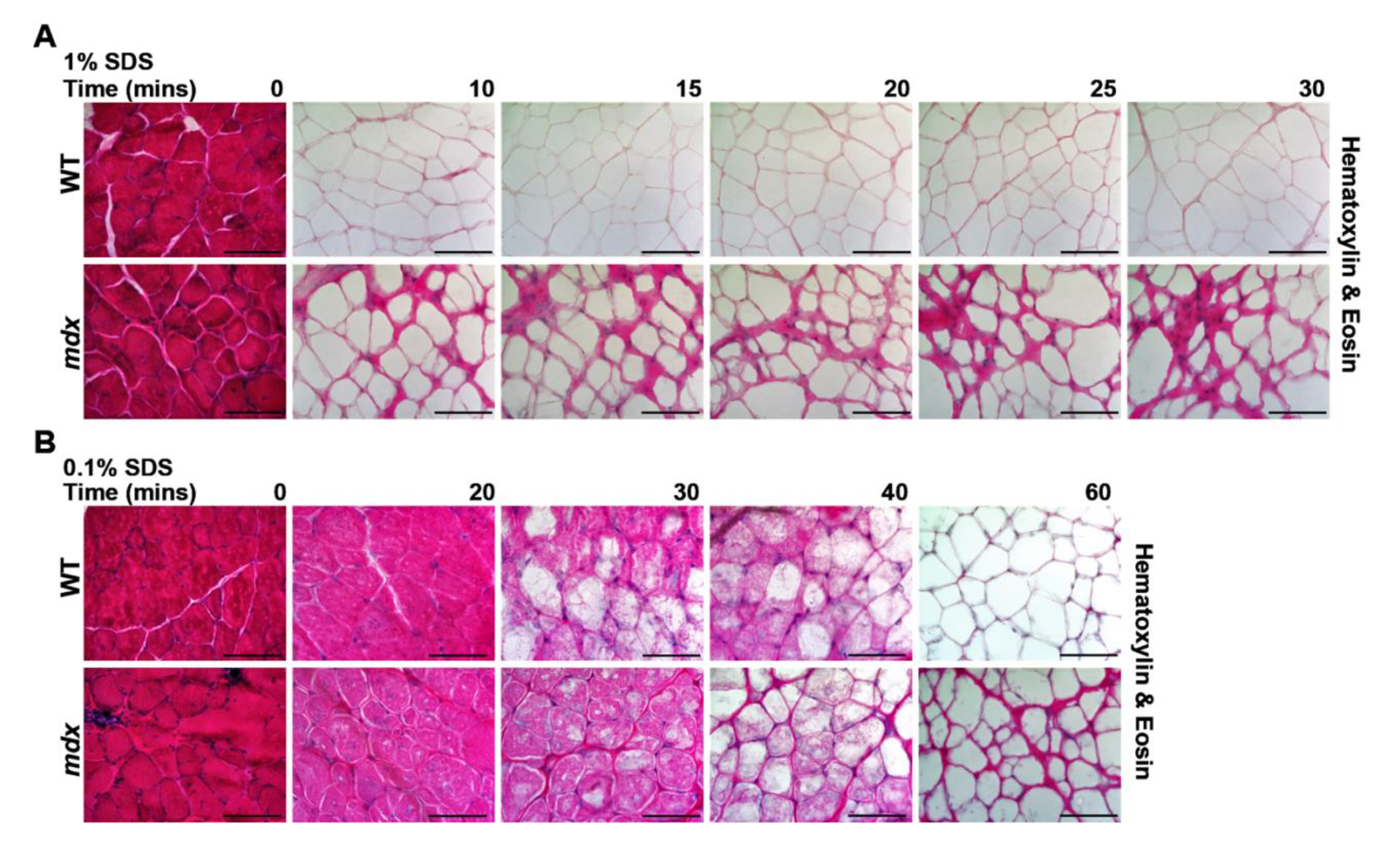
SDS optimization for on-slide decellularization of skeletal muscle. Representative images of Hematoxylin & Eosin (H&E) staining of wild-type (WT) and *mdx* quadriceps muscles that were decellularized with 1% SDS or 0.1% SDS for 10 to 60 minutes. Non-decellularized sections, treated with PBS only, served as controls (0 minutes). (**A**) Exposure of slides to 1% SDS solution produced complete decellularization of the myoscaffolds after 10 minutes. (**B**) At 0.1% SDS, longer incubation times were needed for decellularization, with loss of visualized cellular components achieved at 60 minutes. Scale bar, 100μm.

### Acellular myoscaffolds retain key ECM proteins and architectural integrity

We interrogated the protein composition of dECMs from multiple muscular dystrophy mouse models including mice lacking dystrophin (*mdx*), sarcoglycan (*Sgcg*), or dysferlin (*BLAJ*). We also evaluated the effect of genetic background by including dystrophic models on the DBA/2J (D2) background (*mdxD2* and *SgcgD2*) since this background induces more intense fibrosis (**Supplemental Table 1**). The mutations underlying *mdx* and *Sgcg* mice disrupt sarcolemma stability, while the *BLAJ* model is defective in membrane repair. In each dECM model, matrix quality was assessed following decellularization to determine retention of key proteins and architectural integrity (Crapo et al., 2011; Gilpin and Yang, 2017). Complete removal of the cellular components in the dECMs across all strains was confirmed by H&E staining (**Figure 2A**), and Sirius Red staining evaluated collagen deposition and integrity (**Figure 2B**). Intact collagen structure was visualized within the dECMs with increased collagen deposition observed in more severe muscular dystrophy models as expected based on disease severity (*mdxD2* > *mdx* > WT). Retention of glycosaminoglycans (GAGs) post decellularization was confirmed with Alcian Blue staining (**Figure 2C**).

**Figure 2.**
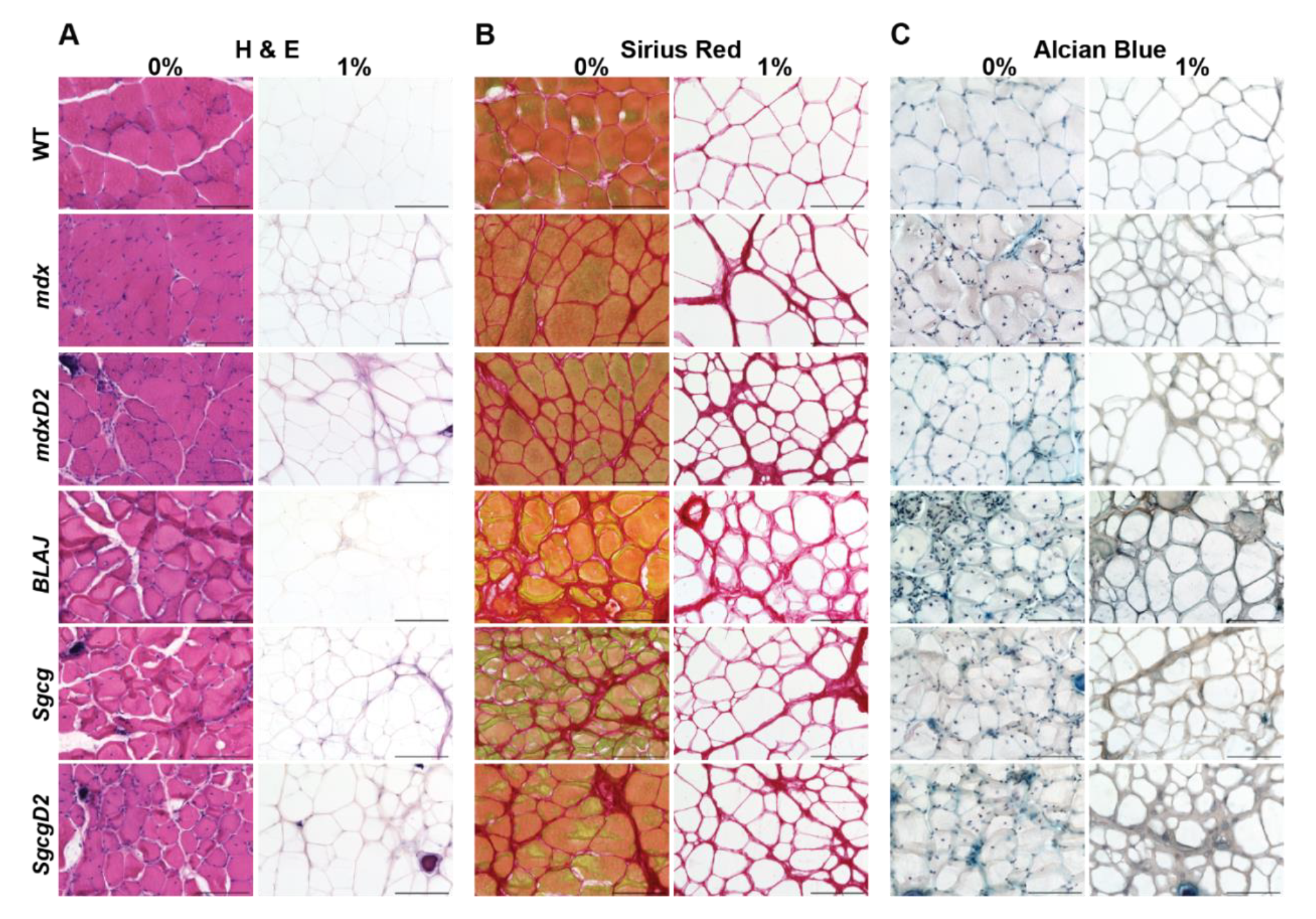
Acellular myoscaffolds retained ECM architectural integrity and glycosaminoglycans after on-slide decellularization. Muscle sections from WT and multiple muscular dystrophy mouse models including *mdx*, *mdxD2*, *BLAJ*, *Sgcg*, and *SgcgD2* were decellularized with a 1% SDS solution (1%) for 10 minutes. Details for each mouse strain are shown in Supplemental Table 1. The non-decellularized control was incubated in PBS (0%). Representative images of (**A**) H&E, (**B**) Sirius Red, and (**C**) Alcian Blue staining demonstrate removal of visualized cellular components, preservation of the dECM collagen structure and retention of glycosaminoglycans respectively. Scale bar, 100μm.

The architectural integrity of the acellular myoscaffolds was further interrogated through immunofluorescence microscopy (IFM) of laminin and collagen pre-and post-decellularization. Antibodies against laminin α2 (LAMA2) and collagen type 1 demonstrated intact laminin and collagen within the dECM scaffolds across all strains (**Figure 3A-B**). Interestingly, both ECM proteins were more readily visualized in acellular scaffolds. This pattern is thought to reflect enhanced epitope exposure after cellular removal (Baptista et al., 2011; Hussein et al., 2016; Stearns-Reider *et al*., 2023). Elimination of nucleic acids was confirmed using Hoechst staining (**Figure 3C**). Additionally, we confirmed the SDS method reduced DNA remnants as expected by comparing DNA content from decellularized and non-decellularized tissue sections. Exposure to 1% SDS decellularization reduced intact DNA to less than 50ng dsDNA per mg ECM (**Supplemental Figure 1**). These data demonstrate that a 10-minute incubation with 1% SDS resulted in intact dECMs that are devoid of cellular components and nucleic acids while retaining core components of ECM maintaining architectural integrity.

**Figure 3.**
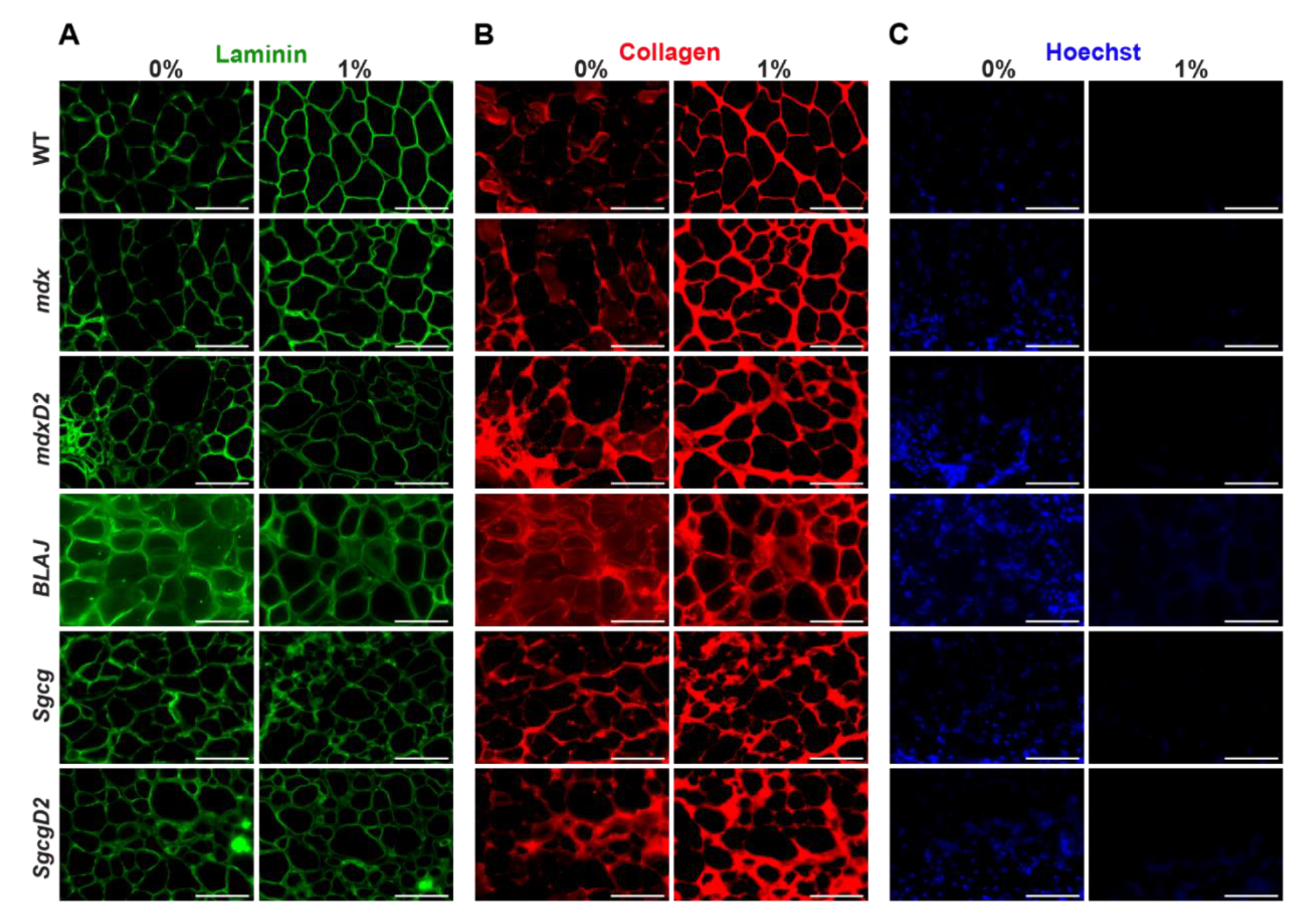
Acellular myoscaffolds from dystrophic models demonstrated excess collagen deposition in all models. Shown is immunofluorescence microscopy (IFM) of 1% SDS decellularized myoscaffolds from WT and muscular dystrophy mouse models (*mdx*, *mdxD2*, *BLAJ*, *Sgcg*, and *SgcgD2*). PBS was used for a non-decellularized control (0%). Representative images of (**A**) laminin-2 (α-2 chain) (**B**) collagen I, and (**C**) Hoechst staining demonstrate the intact nature of the ECM protein. All dECMs displayed brighter collagen signal, consistent with detergent-enhanced epitope unmasking. Greater collagen detection in the muscular dystrophy models was evident with broader regions of collagen deposition. Hoechst staining confirms removal of cellular DNA. Scale bar, 100μm.

### Core matrisome proteins are increased in dystrophin-deficient dECM while only periostin is increased in dysferlin-deficient dECM

ECM components can be separated into “core” and “associated” matrisome proteins (Naba et al., 2016; Shao et al., 2019). Core matrisome proteins include nearly 275 proteins which fall into three categories: glycoproteins (194), collagens (44), and proteoglycans (36) (**Figure 4A**) (Naba *et al*., 2016; Shao *et al*., 2019). We evaluated several core matrisome proteins including decorin (DCN), periostin (POSTN), and thrombospondin-4 (THBS4) since each of these proteins has been implicated in the muscular dystrophy process (Caceres et al., 2000; Chen et al., 2000; Fadic et al., 2006; Lorts et al., 2012; Vanhoutte et al., 2016; Zanotti et al., 2005) (**Figure 4B**). Antibodies to DCN, POSTN, and THBS4 each displayed increased signal in the dECMs from the more severely fibrotic muscular dystrophy muscles (**Figure 4C**). Excess DCN, POSTN, and THBS4 were each observed in the myoscaffolds of *mdxD2* compared to *mdx* and increased in *mdx* compared to WT (**Figure 4C**).

**Figure 4.**
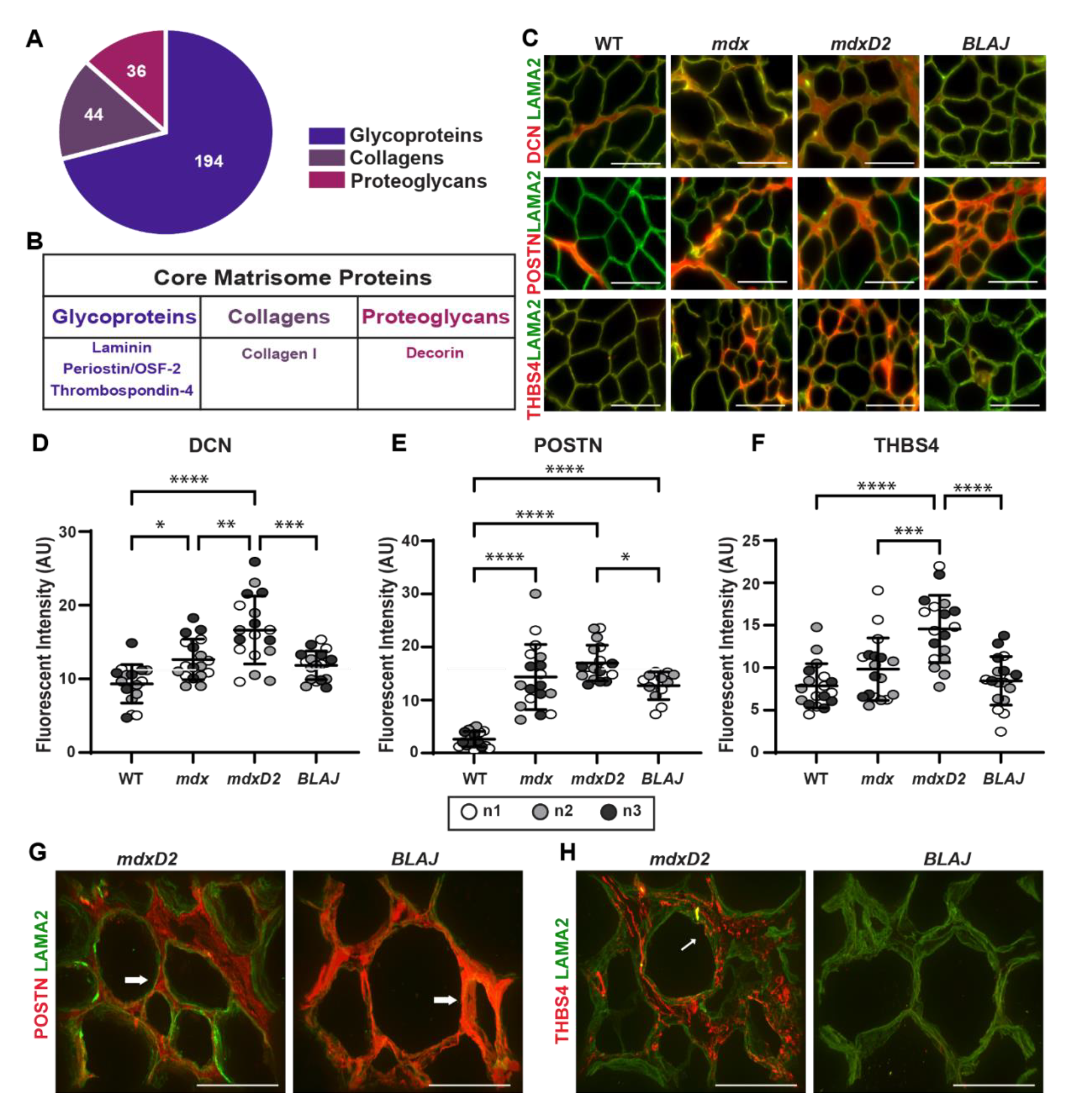
Acellular myoscaffolds differentially retained core matrisome proteins decorin, periostin and thrombospondin 4 in different muscular dystrophy models. (**A**) Core matrisome protein distribution as defined by (Naba *et al*., 2012). (**B**) Proteins selected for analysis in dECMs. (**C**) Representative IFM images from WT and *mdx*, *mdxD2*, and *BLAJ* dECMs showing co-staining with antibodies against decorin (DCN), periostin (POSTN), or thrombospondin-4 (THBS4) (red) with laminin-2 (α-2 chain) (LAMA2, green). DCN was observed diffusely throughout dystrophic dECMs from the DBA/2J background (*mdxD2*). POSTN was increased in all muscular dystrophy subtypes. THBS4 protein expression, like DCN, was more focally upregulated in specific regions of the dystrophic myoscaffolds and was especially increased in muscular dystrophy models in the DBA/2J background. Scale bar, 100μm. Quantitation of IFM signal demonstrated a significant increase in mean fluorescent intensity in myoscaffolds (**D**) DCN (WT 9.7, *mdx* 13.2, *mdxD2* 17.3, *BLAJ* 12.3 AU), (**E**) POSTN (WT 2.6, *mdx* 14.3, *mdxD2* 17.0, *BLAJ* 12.7 AU), and (**F**) THBS4 (WT 7.8, *mdx* 9.8, *mdxD2* 14.6, *BLAJ* 8.5 AU). n=3 independent mice per strain (marked as n1, n2, n3), n=6 images from 2 scaffolds per mouse. (**G,H**) 100x Z-stack representative images of POSTN or THBS4 (red) with LAMA2 (green) demonstrate unique protein deposition profiles on the acellular myoscaffolds. POSTN was increased in dysferlin-deficient *BLAJ*, nearly obliterating the LAMA2 signal (right). In contrast, POSTN signal was interspersed with LAMA2 signal in *mdxD2* dECMs (left). In *mdxD2* dECM, THBS4 was observed in the matrix in a vesicle-like pattern (thin white arrow). Scale bar, 50μm. Graphs show the mean with SEM bars. * p < 0.05, ** p < 0.01, *** p < 0.005, and **** p < 0.001 by one-way ANOVA.

Quantitative analysis of the fluorescence patterns of DCN, POSTN, and THBS4 protein expression in acellular myoscaffolds from WT, *mdx*, *mdxD2*, and *BLAJ* is shown in **Figure 4D-F**. DCN, POSTN, and THBS4 intensity was significantly increased in *mdxD2* acellular dECM myoscaffolds compared to *mdx* and WT matrices. Fluorescence intensity was also increased in *mdx* compared to WT for DCN and POSTN. A consistent pattern of *mdxD2 > mdx > WT* was observed (**Figure 4D-F**). Acellular myoscaffolds generated from the *BLAJ* muscle had significantly less matrix protein compared to *mdxD2* for all three core matrisome proteins analyzed, with protein levels trending similarly with the *mdx* scaffolds for POSTN and DCN (**Figure 4D and E**). Interestingly, THBS4 levels in *BLAJ* dECMs were similar to WT THBS4 levels (**Figure 4F**). WT myoscaffolds generated from the DBA/2J strain had similar expression patterns as WTB6 with minimal DCN, POSTN, THBS4 signal, indicating the dystrophic process contributed to pathological matrix remodeling (**Supplemental Figure 2A**). Immunoblotting of soluble and insoluble muscle fractions isolated from WT, *mdx*, *mdxD2*, and *BLAJ* mice for DCN, POSTN, THBS4 mirrored the IFM quantitation (**Supplemental Figure 3**).

Dysferlin-deficient *BLAJ* dECMs had a distinct pattern of dECM remodeling with only POSTN showing an increased signal compared to WT. This increased POSTN pattern in *BLAJ* dECMs nearly obliterated laminin α2 (LAMA2) staining, in contrast to the pattern seen in *mdxD2* dECMs, where laminin α2 regions were interspersed with POSTN around myofibers (**Figure 4G**). The lack of increased THBS4 was confirmed in dysferlin-deficient *BLAJ* dECMs, and additionally THBS4 had a vesicular pattern in dECMs from *mdxD2* (**Figure 4H**). These data demonstrate that core matrisomal proteins differ across different forms of muscular dystrophy.

### Matrisome-Associated Proteins, MMP9, TIMP-1, and TGF-β1, in acellular myoscaffolds

The matrisome associated protein category consists of over 800 proteins separated into three sub-categories: Secreted Factors (367), ECM-Affiliated (165), and Regulators (304) (**Figure 5A**) (Naba *et al*., 2016; Shao *et al*., 2019). We evaluated muscular dystrophy relevant proteins from each category including annexin A2 (ANXA2), annexin A6 (ANXA6), matrix metalloproteinase-9 (MMP9), TIMP metallopeptidase inhibitor 1 (TIMP-1), and transforming growth factor beta 1 (TGF-β1) (**Figure 5B**). After decellularization, acellular myoscaffolds were probed with antibodies to MMP9, TIMP-1, or TGF-β1 and with anti-laminin α2 (**Figure 5C**). TIMP-1 was distributed uniformly throughout the matrix in all genetic models, while MMP9 and TGF-β1 had distinct focal patterns of distribution in dystrophic dECMs. *mdxD2* had significantly greater MMP9 and TGF-β1 compared to *mdx*, and *mdx* expression levels were greater than WT (**Figure 5D-5F**). DBA/2J WT myoscaffolds demonstrated similar expression patterns as WTB6 with minimal ANXA2, ANXA6, MMP2, TIMP1, and TGF-β1 signal present in the myoscaffold (**Supplemental Figure 2B**). dECM myoscaffolds from dysferlin-deficient *BLAJ* muscle tissues had MMP9 and TGF-β1 protein levels similar to or slightly increased compared to *mdx*, respectively. TIMP-1 expression was increased over WT in a similar fashion across *mdx*, *mdxD2*, and *BLAJ* (**Figure 5D-5F**). Thus, acellular dECM myoscaffolds retained key core and matrisome-associated proteins and these patterns differed across different muscular dystrophy subtypes.

**Figure 5.**
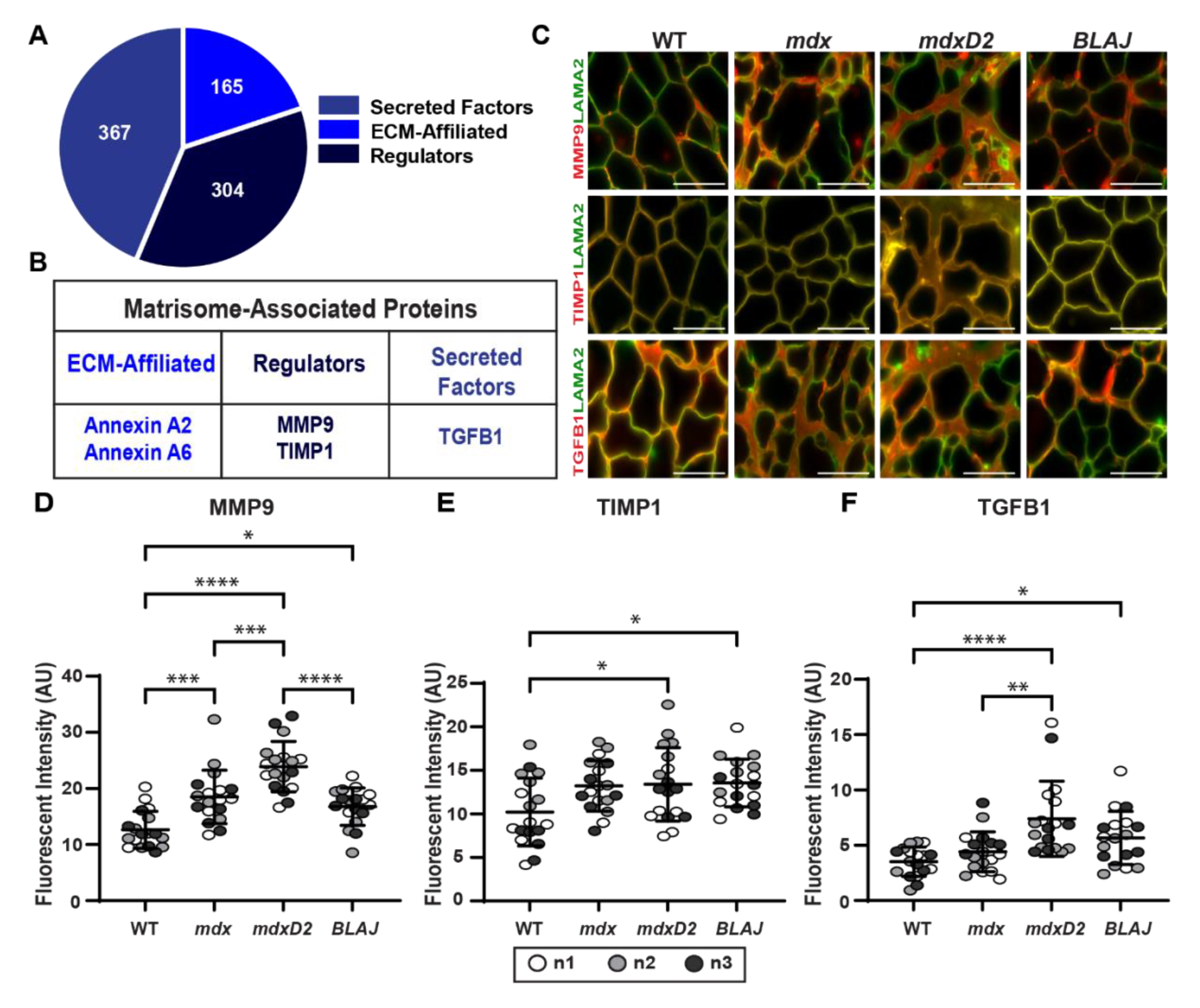
Acellular myoscaffolds retained matrisome-associated proteins MMP9, TIMP-1, and TGFβ-1 and demonstrate differential expression across different dECM models. (**A**) Matrisome-associated protein distribution as defined by (Naba *et al*., 2012). (**B**) Selected proteins studied in dECMs. (**C**) Representative IFM images of dECMs co-stained with antibodies to laminin-2 (α-2 chain) (LAMA2, green) and MMP9, TIMP-1, or TGF-β1 (red). MMP9 and TGF-β1 dECM content was increased in muscular dystrophy models from the DBA/2J background. Scale bar, 100μm. (**D**) MMP9 (WT 12.7, *mdx* 18.5, *mdxD2* 23.9, *BLAJ* 16.8 AU), (**E**) TIMP-1 (WT 10.2, *mdx* 13.2, *mdxD2* 13.4, *BLAJ* 13.6 AU), and (**F**) TGF-β1 (WT 3.5, *mdx* 4.4, *mdxD2* 7.4, *BLAJ* 5.7 AU) mean fluorescent intensity in myoscaffolds. n=3 independent mice per strain (marked as n1, n2, n3), n=6 images from 2 scaffolds per mouse. Graphs show the mean with SEM bars. * p < 0.05, ** p < 0.01, *** p < 0.005, and **** p < 0.001 by one-way ANOVA.

### Annexins A2 and A6 were significantly increased in BLAJ dECM compared to other muscular dystrophy myoscaffolds

The annexin protein family members, including annexin A2 and A6, belong to the matrisome-affiliated category in the subcategory of ECM-affiliated proteins (Naba *et al*., 2016; Naba et al., 2012; Shao *et al*., 2019). Acellular dECM myoscaffolds were probed with antibodies to ANXA2 or ANXA6 and anti-laminin α2 (LAMA2) demonstrating increased ANXA2 and ANXA6 in the matrix of more severely fibrotic models of muscular dystrophy including *mdxD2* and *SgcgD2* (**Figure 6A; Supplemental Figure 4**). Quantitative fluorescence intensity analysis indicated that ANXA2 and ANXA6 signal in *mdxD2* acellular dECM myoscaffolds was increased compared to WT, as expected (**Figure 6A and 6B**). Interestingly, the *BLAJ* dECM had markedly greater ANXA2 and ANXA6 signal compared to all other strains (**Figure 6A, B and C**). The two annexins displayed different expression profiles with ANXA2 exhibiting lower-level, diffuse expression within the matrix compared to ANXA6, which was abundant in vesicle-like structures (**Figure 6D**). Z-stack images showed the increased accumulation of ANXA6 vesicles especially in the dECM of *BLAJ* scaffolds (**Figure 6E**; thin white arrow). However, in areas of high ANXA6 deposition, ANXA6 appeared as a solid rim over laminin-α2, surrounding what remained of the acellular myofibers (**Figure 6E**; thick white arrow). To investigate the origin of matrix-deposited ANXA6 we generated human GFP-tagged ANXA6 utilizing an adeno-associated virus 9 (AAV9) vector driven by the striated muscle specific promotor: MHCK7 (AAV9-MHCK7-ANXA6-GFP; **Supplemental Figure 5**). Mice injected with AAV9-MHCK7-ANXA6-GFP exhibited ANXA6-GFP expression in quadriceps muscle sections compared to no visable signal in PBS injected mice (**Figure 6F**). Acellular myoscaffolds generated from AAV9-injected quadriceps sections displayed ANXA6-GFP deposition in the ECM (**Figure 6G**; white arrows) compared to dECMs generated from PBS-injected control mice, indicating that the observed ANXA6 originates from myofibers. These data demonstrate that ANXA6 deposition into the ECM from muscle cells differs across genotypes with dysferlin-deficient muscular dystrophy having excess annexin deposition in its matrix.

**Figure 6.**
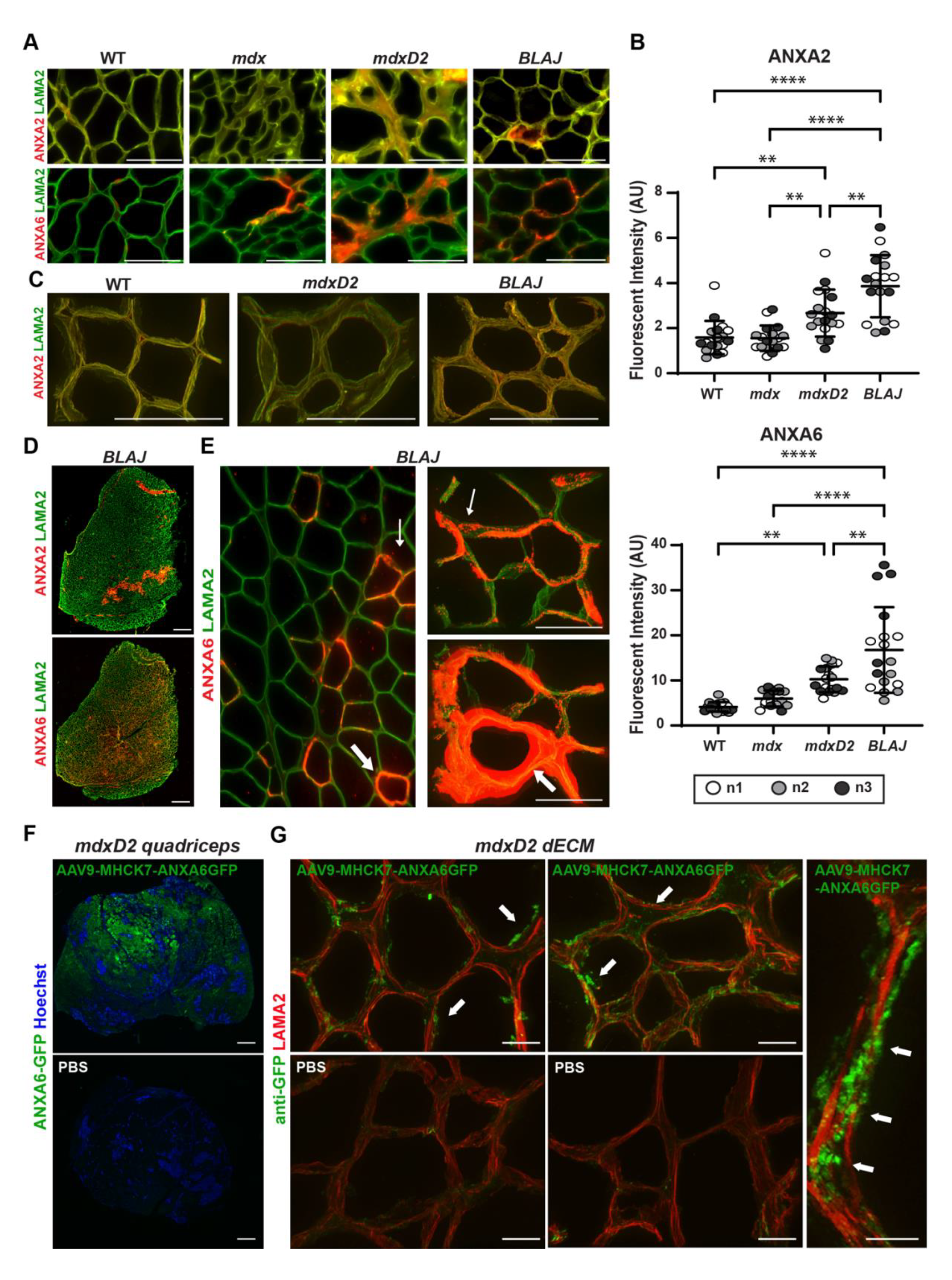
Excess deposition of annexins in dysferlin deficient myoscaffolds. (**A**) Representative IFM images from acellular myoscaffolds generated from WT and *mdx*, *mdxD2*, and *BLAJ* muscles evaluated for annexin A2 (ANXA2) and annexin A6 (ANXA6) (red) with anti-laminin-2 (α-2 chain) (LAMA2, green). ANXA2 exhibits a diffuse pattern, while ANXA6 is vesicular. All scale bars are set at 100µm. (**B**) ANXA2 (WT 1.6, *mdx* 1.6, *mdxD2* 2.7, *BLAJ* 3.9 AU) and ANXA6 (WT 4.1, *mdx* 6.0, *mdxD2* 10.2, *BLAJ* 16.8 AU) mean fluorescent intensity in myoscaffolds was significantly upregulated in *BLAJ* scaffolds. n=3 independent mice per strain (marked as n1, n2, n3), n=6 images from 2 scaffolds per mouse. (**C**) 100x Z-stack images of ANXA2 (red) with LAMA2 co-staining (green) demonstrated diffuse expression on the dECMs generated from WT, *mdxD2* and *BLAJ* muscle. Scale bars 100µm. (**D**) Tiled image of a full thickness *BLAJ* dECM scaffold. ANXA6 staining was present throughout the myoscaffold, while ANXA2 staining was concentrated into focal areas. Scale bars 500µm. (**E**) 20x and 100x Z-stack images of ANXA6 illustrated a unique protein deposition profile on *BLAJ* myoscaffolds. ANXA6 (red) appeared in the matrix in a vesicle-like pattern (thin white arrow), which formed a concentrated ring of protein (thick white arrows). Scale bars 100µm. (**F**) GFP fluorescent imaging revealed robust ANXA6-GFP expression (green) in quadriceps muscle from *mdxD2* mice systemically injected with adeno-associated virus 9 (AAV9) driven by the MHCK7 chimeric striated muscle specific promoter, while a lack of GFP signal was noted in PBS treated muscle. (**G**) Acellular myoscaffolds were generated from AAV9-MHCK7-ANXA6-GFP and PBS treated mice. Z-stack immunofluorescent imaging revealed anti-GFP vesicular staining (green), representing the presence of muscle derived ANXA6-GFP, within the matrix of AAV9 muscle with minimal signal in PBS treated muscle. LAMA2 (red) staining marks the matrix. Scale 20µm low magnification and 10µm high magnification images. Graphs show the mean with SEM bars. * p < 0.05, ** p < 0.01, *** p < 0.005, and **** p < 0.001 by one-way ANOVA.

### Dystrophic dECMs adversely impact migratory parameters of C2C12 myoblasts seeded onto myoscaffolds

Muscle regeneration is generally impaired in severely dystrophic muscle (Wallace and McNally, 2009), with *mdxD2* mice having even more impaired muscle regeneration compared to *mdx* muscle (Coley *et al*., 2016). Stearns-Reider showed that skeletal muscle progenitor cells derived from human pluripotent stem cells had reduced mobility and differentiation on *mdx* dECMs (Stearns-Reider *et al*., 2023). We evaluated the migration patterns of undifferentiated C2C12 myoblasts seeded onto dECMs prepared from multiple dystrophic models (**Figure 7A**). C2C12 myoblasts were allowed to seed for 24 hours then, cell migration was recorded for the next 24-48 hours with images acquired every 20 minutes (experimental timeline shown in **Supplemental Figure 6**). C2C12 myoblasts seeded on *mdx* and *mdxD2* dECMs had reduced movement throughout the matrix compared to *BLAJ* and *WT* dECMs (**Figure 7B**; black arrows; movement quantified in **Figure 7C**). Conversely, an increase in the frame-frame distance was seen for C2C12 myoblasts seeded on *BLAJ* compared to *WT* dECMs (**Figure 7C**). A similar pattern was observed in the distanced traveled from the start position by C2C12 myoblasts on the matrices with *BLAJ* > *WT* > *mdx* > *mdxD*2 (**Figure 7D**). Myoblast speed was also affected by the dECM substrate. Myoblasts seeded on *BLAJ* dECMs demonstrated greater speed compared to *WT*, *mdx* and *mdxD2* (**Figure 7E**), while myoblasts seeded on *mdx* and *mdxD2* myoscaffolds had reduced speed compared to WT scaffolds (**Supplemental video 1 and 2)**.

**Figure 7.**
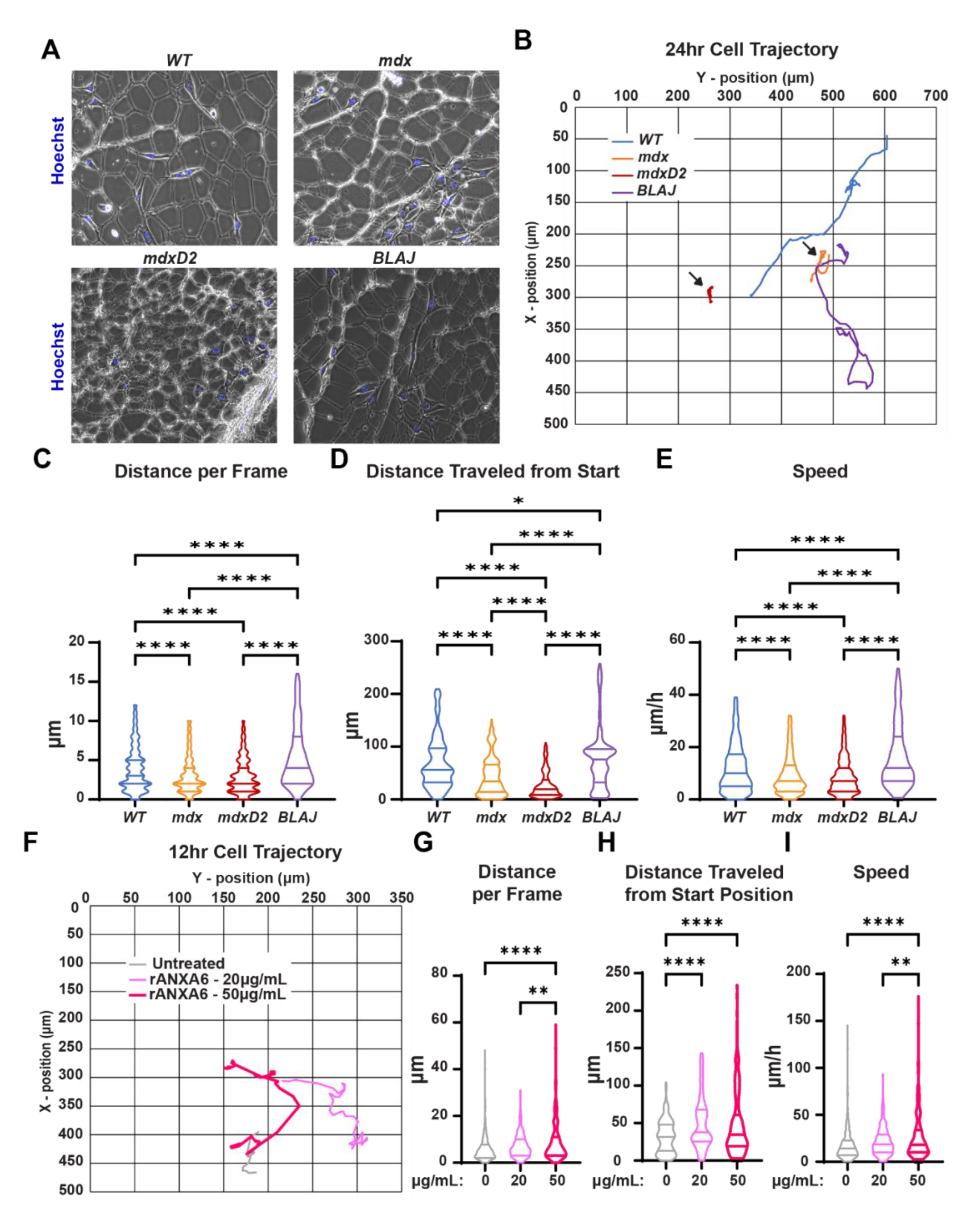
*mdx* myoscaffolds inhibit C2C12 myoblasts movement while *BLAJ* myoscaffolds enhance C2C12 myoblast movement. (**A**) Representative phase contrast images of C2C12 myoblasts seeded onto dECMs from WT, *mdx*, *mdxD2*, and *BLAJ mice*. (**B**) Blue (WT), orange (*mdx*), red (*mdxD2*), and purple (*BLAJ*) lines depict cellular movement trajectory over 24 hours. Quantification of C2C12 cell movement demonstrated a significant difference in (**C**) distance per frame (*WT* 3.91, *mdx* 3.10, *mdxD2* 2.98, and *BLAJ* 5.41 μm), (**D**) distance traveled from start position (*WT* 69.24, *mdx* 43.99, *mdxD2* 27.12, and *BLAJ* 76.78 μm), and (**E**) speed (*WT* 11.89, *mdx* 9.31, *mdxD2* 8.80, and *BLAJ* 16.34 μm/h) among the WT and dystrophic myoscaffolds. Myoscaffolds prepared from two mice per genotype; n=73 images per cell, with 8-11 myoblasts tracked per genotype. C2C12 cells were treated with recombinant ANXA6 (rANXA6) and tracked; (**F**) Grey (untreated), light pink (20 μg/mL rANXA6) and pink (50 μg/mL rANXA6) lines depict cellular movement trajectory over 12 hours. Quantification of rANXA6-treated myoblasts demonstrated a significant difference in (**G**) distance per frame (0 μg/mL 6.01, 20 μg/mL 7.23, and 50μg/mL 12.09 μm), (**H**) distance traveled from start position (0 μg/mL 32.18, 20 μg/mL 49.72, and 50μg/mL 55.43 μm), and (**I**) speed (0 μg/mL 18.04, 20 μg/mL 21.78, and 50 μg/mL 36.20 μm). Treatments were conducted on two different passages; n=37 images per cell, with 8 myoblasts tracked per treatment. Graphs show the mean with SEM bars. * p < 0.5, ** p < 0.01, *** p < 0.005, and **** p < 0.001 by one-way ANOVA.

Because the altered matrix composition observed in dysferlin-deficient matrices was characterized by increased ANXA6 deposition, and this same matrix supported increased myoblast motility, we assessed whether ANXA6 itself contributed to differential cellular behavior. C2C12 myoblasts cultured on standard plates were treated with 0, 20, or 50μg/mL of recombinant ANXA6 (rANXA6) and cell migration was recorded for 12-24 hours with images acquired every 20 minutes. C2C12 myoblast cells treated with rANXA6 had a dose-dependent increase in movement per frame (**Figure 7F**; movement per frame quantified in **Figure 7G**; **Supplemental video 3 and 4**). A similar pattern was observed in the distanced traveled from the start position by rANXA6 treated C2C12 myoblasts with 50 μg/mL > 20 μg/mL > 0 μg/mL (**Figure 7H**). Myoblast speed was also affected by the rANXA6 with increasing dosage resulting in increased speed (**Figure 7E**). Combined, these data indicate that disease-specific matrix composition results in unique cellular cross-talk directly altering cellular behavior and that ANXA6 itself can contribute to the increase in myoblast mobility.

## DISCUSSION

The process of tissue decellularization has been widely applied in regenerative medicine as a means of testing biological signaling properties that support recellularization. The adaptation of this method to an “on-slide” format for dystrophic muscle enables an evaluation of how the ECM differs across muscular dystrophy subtypes (Stearns-Reider *et al*., 2023). On-slide dECM studies preserve the spatial architecture and composition of the matrix, with decellularization resulting in increased epitope exposure from removing the cellular component, allowing greater ECM protein accessibility and visualization. Stearns-Reider et al. observed that intensely fibrotic regions of *mdx* scaffolds had differential collagen crosslinking and laminin deposition which impacted skeletal muscle progenitor cells (SMPCs) motility and the ability of SMPCs to remodel severe fibrosis. Our findings of impaired C2C12 mobility on severe *mdxD2* scaffolds compared to less fibrotic disease models are consistent with the observations of Stearn-Reider illustrating impaired motility on the more intense areas of fibrosis within *mdxB10* dECMs. Our findings suggest that C2C12 similarly communicate with the ECM impacting function.

We evaluated three different primary mutations that cause muscular dystrophy, and compared how these mutations shifted the myoscaffold content. Two of the primary mutations, dystrophin and γ-sarcoglycan, share a similar pathological defect, namely a fragile sarcolemma that undergoes repetitive disruption and leak. However, dystrophin itself contributes directly to the stiffness of the sarcolemma, while the sarcogclyan complex is implicated in the attachment of sarcolemma to the surrounding matrix. Thus, the finding that the dECMs similarly impact matrix remodeling supports the defect in sarcolemmal stability as the major contributor to fibrosis. Both *mdx* and *Sgcg* mutants shared myoscaffold features, including similar matrisome protein profiles and associated fibrosis, and these features were even more evident in the DBA/2J background, which is known to produce more fibrosis (Coley *et al*., 2016; Hammers *et al*., 2020; Heydemann *et al*., 2005). In particular, these models, *mdx* and *Sgcg,* in the DBA/2J background had significantly increased levels of core matrisomal proteins THBS4, POSTN, and DCN within the dECM scaffolds. THBS4 regulates muscle attachment at the myotendinous junction and promotes sarcolemmal stability (Subramanian et al., 2007; Vanhoutte *et al*., 2016). THBS4 is upregulated in human muscular dystrophies and their corresponding mouse models (Brody et al., 2018; Chen *et al*., 2000; Subramanian and Schilling, 2014; Vanhoutte *et al*., 2016). Overexpression of THBS4 in muscle using transgenesis was shown to protect against muscular dystrophy disease progression by stabilizing the sarcolemma through enhanced trafficking of the dystrophin-glycoprotein and integrin attachment complexes (Brody *et al*., 2018; Vanhoutte *et al*., 2016). Moreover, secretion of THBS4 was shown to be an essential component to elicit these protective effects. Secretion-deficient THBS4 was associated with an increase in intracellular vesicle accumulation visualized by electron microscopy resulting in reduced trafficking to the membrane and more severe disease. We observed THBS4 in puncta in *mdxD2* myoscaffolds throughout the matrix, and these puncta likely reflect the normal secretion of THBS4 vesicles, supporting a role for THBS4 in transporting matrix-bound molecules.

We also observed upregulation of DCN in *mdx* and *Sgcg* myoscaffolds, and this upregulation was further enhanced by the DBA/2J background. DCN is increased in the matrix of DMD muscle biopsies, likely due to the increased synthesis of DCN by DMD fibroblasts (Caceres *et al*., 2000; Fadic *et al*., 2006; Zanotti *et al*., 2005). Upregulation of DCN in myoblasts was shown to improve muscle regeneration, indicating that DCN’s cell of origin impacts its effect (Li et al., 2007). POSTN is normally expressed at low levels in healthy tissues, but its expression is increased after acute injury and in muscular dystrophies (Hamilton, 2008; Lorts *et al*., 2012). Unlike THBS4, overexpression of POSTN exacerbated muscle disease progression and promoted fibrosis formation in *Sgcd*-null mice, a model of LGMD2F (Lorts *et al*., 2012). In dECMs POSTN expression was increased in the more severe *mdxD2* dECM compared to the mild *mdx* and WT dECMs. We hypothesize that the increased TGF-β levels in this model upregulates *Postn* expression while *Dcn* is upregulated as a compensatory protective mechanism.

### Dysferlin deficient muscle displays a distinct dECM content

Dysferlin is a membrane associated protein implicated in resealing disrupted membranes, in contrast to the dystrophin complex and its role in stabilizing the sarcolemma (Bansal *et al*., 2003; Defour *et al*., 2014). Additionally, patients with *DYSF* mutations often have a distinct clinical course where muscle function is normal in early life unlike other forms of muscular dystrophy (Klinge *et al*., 2010). Similarly, mouse models lacking *Dysf* display a comparatively mild phenotype with certain muscles showing more pathology than others (Demonbreun et al., 2014; Nagy et al., 2017; Terrill et al., 2013). The quadriceps muscles, which were used in this study, often have little histopathology at the age used in this study (Demonbreun *et al*., 2014; Nagy *et al*., 2017; Terrill *et al*., 2013). The pattern of increased annexin deposition and its vesicular pattern throughout the dysferlin-deficient *BLAJ* muscle was only seen in dysferlin-deficiency, and it is possible this pattern arises from the increased recruitment of annexin A6 to the membrane lesion in an attempt to enhance membrane resealing. Through immunofluorescence microscopy and correlative light and electron microscopy (CLEM), annexin A6-positive vesicles have been visualized emerging from the membrane lesion seconds to minutes post rupture (Croissant *et al*., 2020; Demonbreun *et al*., 2019a). The annexin vesicles identified in dECM scaffolds are plausibly derived from vesicles released from the injured myofiber repair caps, consistent with our finding that muscle-derived annexin deposits into the decellularized matrix. The distinct localization between vesicular annexin A6 and diffuse annexin A2 in the dECM scaffolds suggests the possibility that these matrix-associated proteins may have different cellular origins. Immunofluorescence imaging of 12-month-old BLAJ muscles showed annexin A2 accumulation in the interstitial space which correlates with fibroblast markers PDGFRa and perilipin-1 (Hogarth et al., 2019).

Additionally, compared to the *mdxD2* matrices, the *BLAJ* dECMs contained less fibrosis, with laminin and collagen content visually similar to WT matrices. Furthermore, myoblasts seeded on the dysferlin-deficient myoscaffolds demonstrated significantly increased cell movement and migration compared to WT, treating myoblasts with recombinant annexin A6 was sufficient to induce increased cellular movement. These findings together provide a possible molecular contribution to the clinical observation *DYSF* deficient patients have preserved or enhanced exercise capacity, and together this work highlights the importance of matrix contributions to muscle health.

## MATERIALS AND METHODS

### Animals

All animals were bred and housed in a pathogen free facility. Wild-type (WT) mice were from the C57BL/6 strain. C57BL/10ScSn-*Dmdmdx/*J (*mdx*), D2.B10-*Dmdmdx/*J (*mdxD2*), and B6.A-*Dysfprmd/*GeneJ (*BLAJ*) mice were originally purchased from The Jackson Laboratory (strains #:001801, #:013141, and #:012767 respectively). Sarcoglycan γ-null (*Sgcg*) mice were previously generated on the C57BL/6 background and Sgcg 521ΔT (*SgcgD2*) were previously generated on the DBA/2J background (Demonbreun et al., 2019b; Hack *et al*., 1998). Seven-to eight-month male and female mice were used for all experiments. Mice were bred and housed in a specific pathogen free facility on a 12-hour light/dark cycle and fed *ad libitum*.

### Study Approval

All procedures using mice followed *Guide for the Care and Use of Laboratory Animals* and were approved by Northwestern University’s Institutional Animal Care and Use Committee.

### Decellularization

Quadriceps muscle was collected and flash frozen in liquid nitrogen. Sections were cryosectioned at 25μm (Leica CM1950) and mounted on charged Superfrost Plus Microscope Slides (cat# 1255015; Fisher Scientific). Slides were stored at -80°C. Decellularization was performed as in Stearns-Reider et al.,2022 with the following modifications (Stearns-Reider et al., 2022). Slides were allowed to thaw to room temperature for 1 hr prior to decellularization. Slides were then placed in a 15mL UltraPure Sodium Dodecyl Sulfate (SDS) solution (cat# 15553035; Invitrogen) diluted with PBS to 0.1 or 1% concentration for 10 minutes under constant agitation (40 rpm). Slides were then placed in 100mm x 100mm square petri dishes 3 slides per dish (Fisher cat#FB0875711A) containing 13mL of PBS (with calcium & magnesium [Cat# 21030CV; Corning]), for 15 minutes, then placed in a fresh 13mL of PBS for 45 minutes, followed by 13mL UltraPure Distilled H_2_O (cat# 10977015; Invitrogen) for 30 minutes, and a final 13mL PBS wash for 45 minutes. All wash steps were done under constant agitation (40 rpm). Decellularized slides were either fixed for histology/immunostaining or stored in PBS.

### DNA Extraction

DNA was extracted from quadriceps muscle tissues using the DNeasy Blood & Tissue Kit (cat# 69504; Qiagen) and DNA concentration measured on a NanoDrop 2000 Spectrophotometer (Thermo Scientific). DNA electrophoresis was visualized using a BioSpectrum Imaging system Chemi 410 (UVP).

### Histology

Quadriceps muscle was harvested and fixed in 10% formalin. Hematoxylin and eosin (cat# 12013B and 1070C; Newcomer Supply), Sirius Red (cat# 24901250; Polysciences, Inc), or Alcian Blue (cat#A3157; Sigma) staining was performed per manufacturer’s protocol. All images were acquired on the Keyence BZ-X810 microscope with a 40x objective and same exposure settings across all genotypes.

### Protein isolation

*Tibialis anterior* muscles were harvested and flash-frozen in liquid nitrogen. Tissues were ground with a mortar and pestle and lysed in Whole Tissue Lysis buffer (50mM HEPES pH 7.5, 150mM NaCl, 2mM EDTA, 10mM NaF, 10mM Na-pyrophosphate, 10% glycerol, 1% Triton X-100, 1mM phenyl-methylsulfonyl fluoride, 1x cOmplete Mini protease inhibitor cocktail [cat # 11836170001; Roche], 1x PhosSTOP [cat# 04906837001; Roche]) and homogenized using a TissueLyser II (Qiagen) at a frequency of 30.0 1/s and time of 2.00 min/sec for 6 rounds. Protein lysates were then centrifuged at 20,000xg for 20 minutes to separate the soluble and insoluble fractions. The soluble supernatant layer was placed in a separate tube and the insoluble pellet layer was resuspended with an 8M Urea buffer (8M Urea [cat#U4883; MilliporeSigma], 2mM EDTA, 10mM DTT, 1x cOmplete Mini protease inhibitor cocktail [cat # 11836170001; Roche]). The protein concentration was measured with the Quick Start Bradford 1x Dye Reagent (cat# 5000205; Bio-Rad Laboratories).

### Immunoblotting

20-25μg of protein lysate with 4x Laemmli buffer (cat# 1610747; Bio-Rad Laboratories was separated on 4-15% Mini-PROTEAN TGX Precast Protein Gels (cat# 4561086; Bio-Rad Laboratories) and transferred to Immun-Blot PVDF Membranes for Protein Blotting (cat# 1620177; Bio-Rad Laboratories). Membranes were reversibly stained for total protein with Ponceau S Staining Solution (cat# A40000278; Thermo Scientific) to ensure transfer and equal loading prior to antibody incubation. Blocking and antibody incubations were done using StartingBlock T20 (TBS) Blocking Buffer (cat# 37543; Thermo Scientific). Primary antibodies were used at 1:1000 except for anti-thrombospondin-4 and anti-TIMP-1 which were diluted at 1:100. Secondary antibodies conjugated with horseradish peroxidase were used at 1:2,500 (cat# 111035003; Jackson ImmunoResearch Laboratories). Clarity Western ECL Substrate (cat# 1705061; BioRad Laboratories) was applies to membranes and visualized using an iBright 1500 Imaging System (Invitrogen). Pierce Reversible Protein Stain Kit for PVDF Membranes (including MemCode; cat# 24585; Thermo Fisher Scientific) was used to ensure equal loading.

### Immunostaining and immunofluorescence imaging

Decellularized quadriceps cryosections (dECMs) were fixed in 4% paraformaldehyde, rinsed, and blocked with a 1% BSA/10% FBS blocking buffer for 1 hour. The dECMs were incubated with the primary antibodies overnight at 4°C at 1:100. The dECMs were then incubated with the secondary antibodies at 1:2,500 for 1 hr. The dECMs were mounted with ProLong Gold Antifade Reagent (cat# P36934; Invitrogen) and images acquired on a Keyence BZ-X810 microscope with 20x, 40x, 60x and 100x objectives. All image exposure settings were kept consistent across genotypes. Fluorescent intensity was quantified using Image J (NIH). Quantification was performed on 18 images per strain, unless otherwise noted. Specifically, 3 representative 40x images were taken from 2 decellularized scaffolds per animal for a total of 6 images per animal.

### Antibodies

Primary antibodies used for immunoblotting and immunostaining: anti-annexin A6 (cat# ab199422; Abcam), anti-annexin A2 (cat# 610068; BD Bioscience), anti-collagen type 1 (cat# CL50151AP1; Cedarlane), anti-decorin (cat# AF1060; R&D Systems), anti-HSP90 (cat#4874; Cell Signaling Technologies), anti-laminin-2 (α-2 Chain) (cat# L0663; MilliporeSigma), anti-MMP9 (cat# ab38898; Abcam), anti-periostin/OSF-2 (cat# NBP130042; Novus Bio), anti-TGF Beta 1 (cat# 218981AP; Proteintech and cat# MA515065; Invitrogen), anti-Thrombospondin-4 (cat# AF2390; R&D Systems), and anti-TIMP-1 (cat# AF980; R&D Systems). Secondary antibodies used: Alexa Fluor 488 goat anti-rat (cat# A11006; Invitrogen), Alexa Fluor 594 goat anti-rabbit (cat# A11012; Invitrogen) and Alexa Fluor 594 donkey anti-goat (cat# A11058; Invitrogen).

### Adeno-associated virus production and dosing

To achieve muscle specific expression of human annexin A6, AAV9-MHCK7-hANXA6-turboGFP was generated and packaged at Vector builder. As a viral plasmid control, liver specific expression of human annexin A6, AAV9-TBG-hANXA6-turboGFP was generated and packaged at Vector builder. At 28days of age, *mdxD2* mice were injected with 5e13 vg/kg AV9-MHCK7-hANXA6-turboGFP or PBS control in a total of 100µl. Four weeks post injection, mice were euthanized. Tissues were harvested and stored at -80C.

### Recombinant protein production

Recombinant human annexin A6 (rANXA6) was generated in *E.Coli* using standard methods (Demonbreun et al., 2022). Protein was diluted in TBS with endotoxin levels less than 1.5 EU/mg.

### Reseeding and live-cell imaging

After decellularization as described above, MatTek chambers (cat# CCS-2; MatTek) were attached to slides mounted with quadricep dECMs. The acellular myoscaffolds were incubated in DMEM media containing 10% FBS and 1% penicillin/streptomycin for 24 hrs prior to reseeding with C2C12 myoblasts (cat# CRL-1772; ATCC). Cells were stained with Hoechst 33342 for 5-minutes and then seeded onto the acellular myoscaffolds at a density of 25,000 in 2mLs media per chamber. Seeded slides were imaged in a temperature and CO2 controlled incubation chamber (STR Stage Top Incubator; Tokai Hit) and images acquired every 20 minutes over 48 hrs on a Keyence BZ-X810 microscope with 20x phase contrast objectives. For recombinant annexin A6 experiments, cells were treated with 0, 20 or 50 µg/ml of recombinant annexin A6 in media. All image exposure settings were kept consistent across all genotypes. Cell tracking analysis was performed using the Dynamic Tracking software on the Keyence.

### Statistical Analyses

Statistical analyses were performed with Prism (GraphPad, La Jolla, CA). Comparisons relied on 1-way ANOVA for 1 variable or 2-way ANOVA for two variables. Otherwise, unpaired two-tailed t-tests were performed. A p-value less than or equal to 0.05 was considered significant. Data were presented as single values were appropriate. Error bars represent ± standard error of the mean (SEM).

## Abbreviations

(ECM): Extracellular matrix
(dECM): decellularized extracellular matrix
(DMD): Duchenne Muscular Dystrophy
(LGMD): Limb Girdle Muscular Dystrophy
(SDS): sodium dodecyl sulfate

## ACKNOWLEDGEMENTS

This work was supported by

National Institutes of Health NS047726 (EM)

National Institutes of Health AR052646 (EM)

National Institutes of Health HL061322 (EM)

National Institutes of Health AR048179 (RC)

National Institutes of Health T32AR065972 (RC)

Additional funding was through Lakeside Discovery (EM, AD).

We especially acknowledge the Jain Foundation for providing dysferlin-null mice from their private colony at The Jackson Laboratory.

## AUTHOR CONTRIBTUITIONS

AL, JK, NR, JO, GL, and AD prepared dECMs and performed immunofluorescence imaging. AL and AD quantified the fluorescence images and performed the migration studies. MH performed mouse husbandry. AL, NR and LV performed the muscle isolations and related immunoblots. AD, LV, and JK performed the AAV study. AD, JK and LV performed the AAV injections and analysis. PP performed histological stains. RC and JR provided dECM methodology and critical input in experimental design. AL, AD and EM conceived and designed the studies, analyzed the data, and wrote the manuscript.

## DATA AND MATERIALS AVAILABILITY

Data available upon request.

---

**SUPPLEMENTAL TABLE 1.**
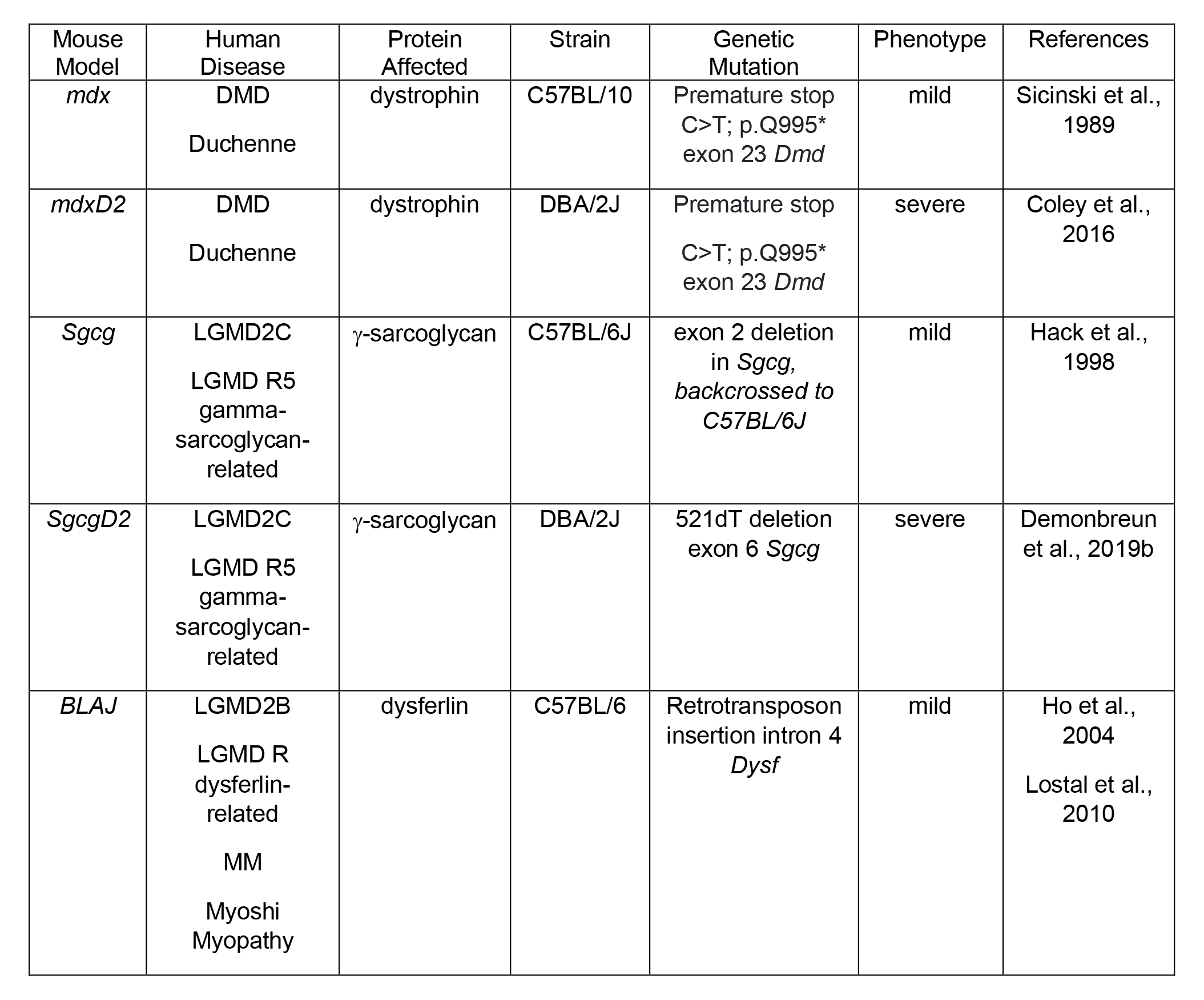
Muscular dystrophy mouse models used in this study.

| Mouse Model | Human Disease | Protein Affected | Strain | Genetic Mutation | Phenotype | References |
| --- | --- | --- | --- | --- | --- | --- |
| *mdx* | DMD<br>Duchenne | dystrophin | C57BL/10 | Premature stop<br>C>T; p.Q995*<br>exon 23 *Dmd* | mild | Sicinski et al.,<br>1989 |
| *mdxD2* | DMD<br>Duchenne | dystrophin | DBA/2J | Premature stop<br>C>T; p.Q995*<br>exon 23 *Dmd* | severe | Coley et al.,<br>2016 |
| *Sgcg* | LGMD2C<br>LGMD R5<br>gamma-<br>sarcoglycan-<br>related | γ-sarcoglycan | C57BL/6J | exon 2 deletion<br>in *Sgcg*,<br>*backcrossed to*<br>*C57BL/6J* | mild | Hack et al.,<br>1998 |
| *SgcgD2* | LGMD2C<br>LGMD R5<br>gamma-<br>sarcoglycan-<br>related | γ-sarcoglycan | DBA/2J | 521dT deletion<br>exon 6 *Sgcg* | severe | Demonbreun<br>et al., 2019b |
| *BLAJ* | LGMD2B<br>LGMD R<br>dysferlin-related<br>MM<br>Myoshi<br>Myopathy | dysferlin | C57BL/6 | Retrotransposon<br>insertion intron 4<br>*Dysf* | mild | Ho et al.,<br>2004<br>Lostal et al.,<br>2010 |

## SUPPLEMENTAL FIGURES AND LEGENDS

**Supplemental Figure 1.**
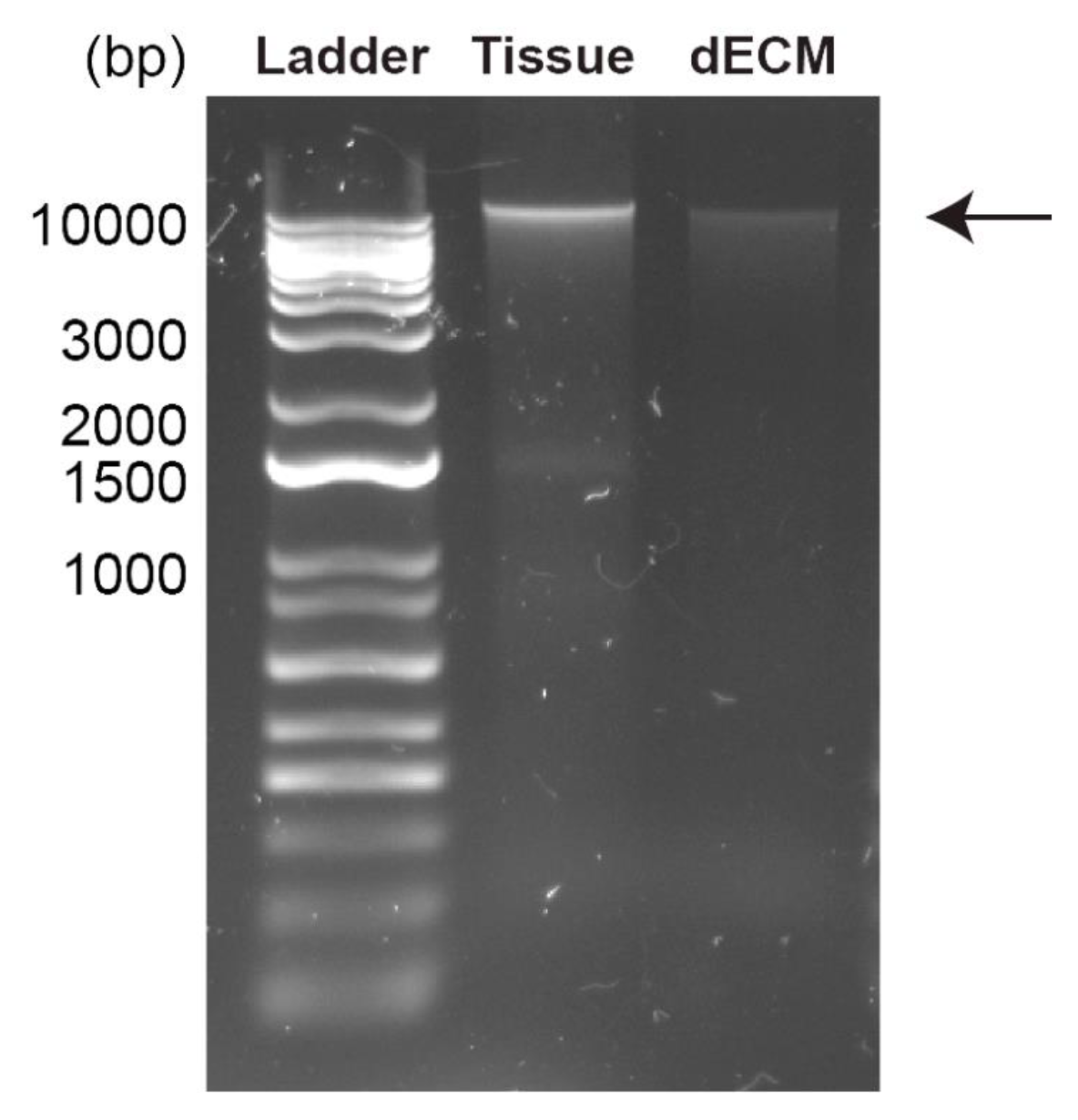
Decreased DNA content after 1% SDS decellularization.

**Supplemental Figure 2.**
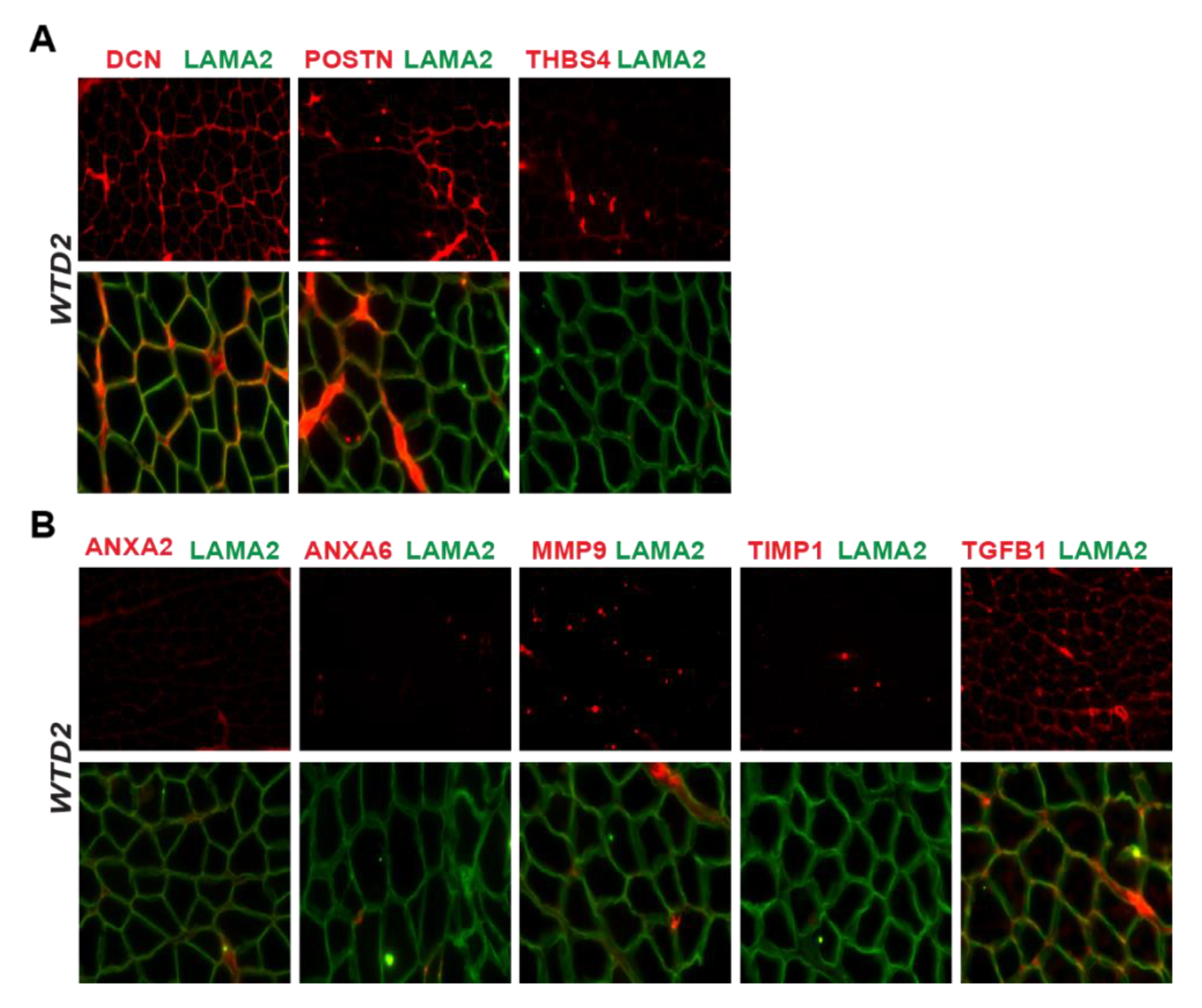
Representative IFM images from acellular myoscaffolds generated from WTD2 mice. (**A**) Core Matrisome Proteins: DCN, POSTN, and THBS4. (**B**) Matrisome-Associated Proteins: ANXA2, ANXA6, MMP9, TIMP1, and TGF-β1.

**Supplemental Figure 3.**
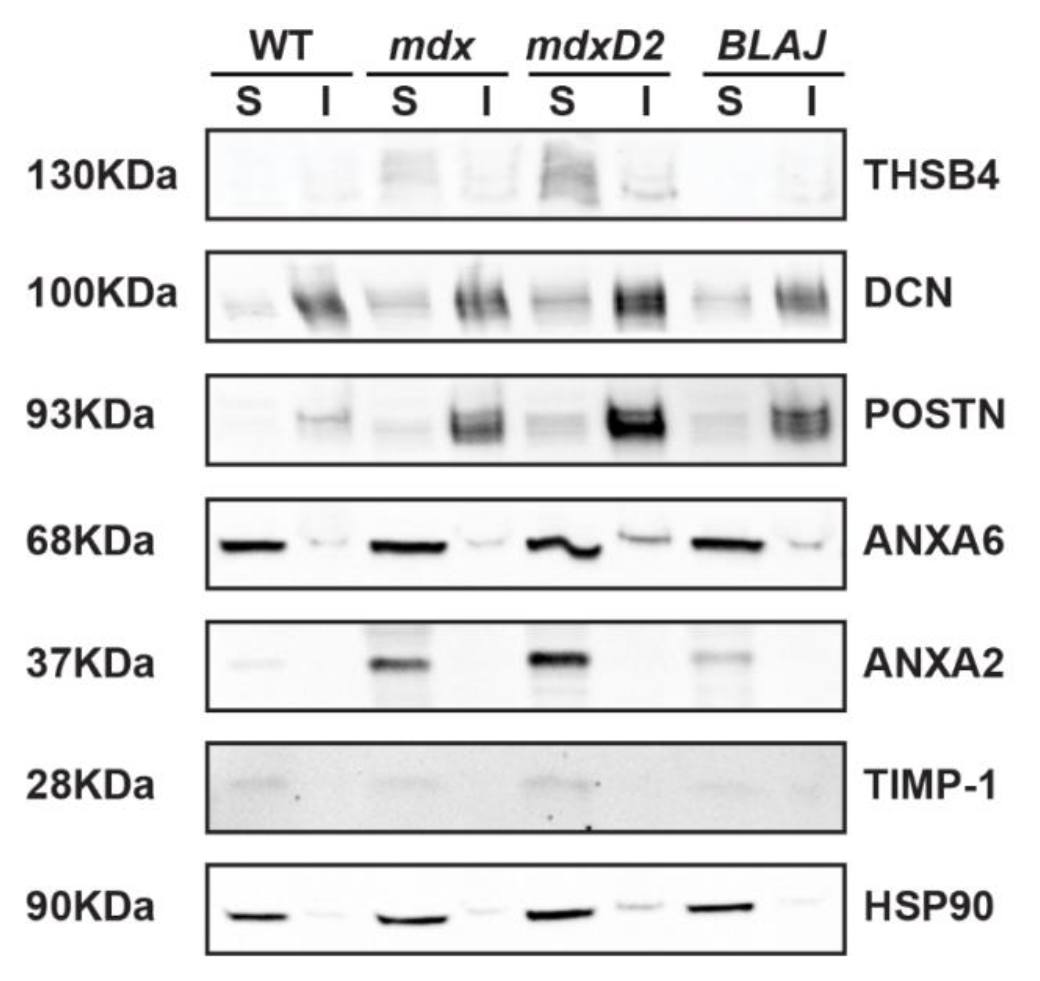
Immunoblot of soluble (S) and insoluble (I) muscle lysate fractions reveals enrichment of matrix associated proteins in dystrophic muscle.

**Supplemental Figure 4.**
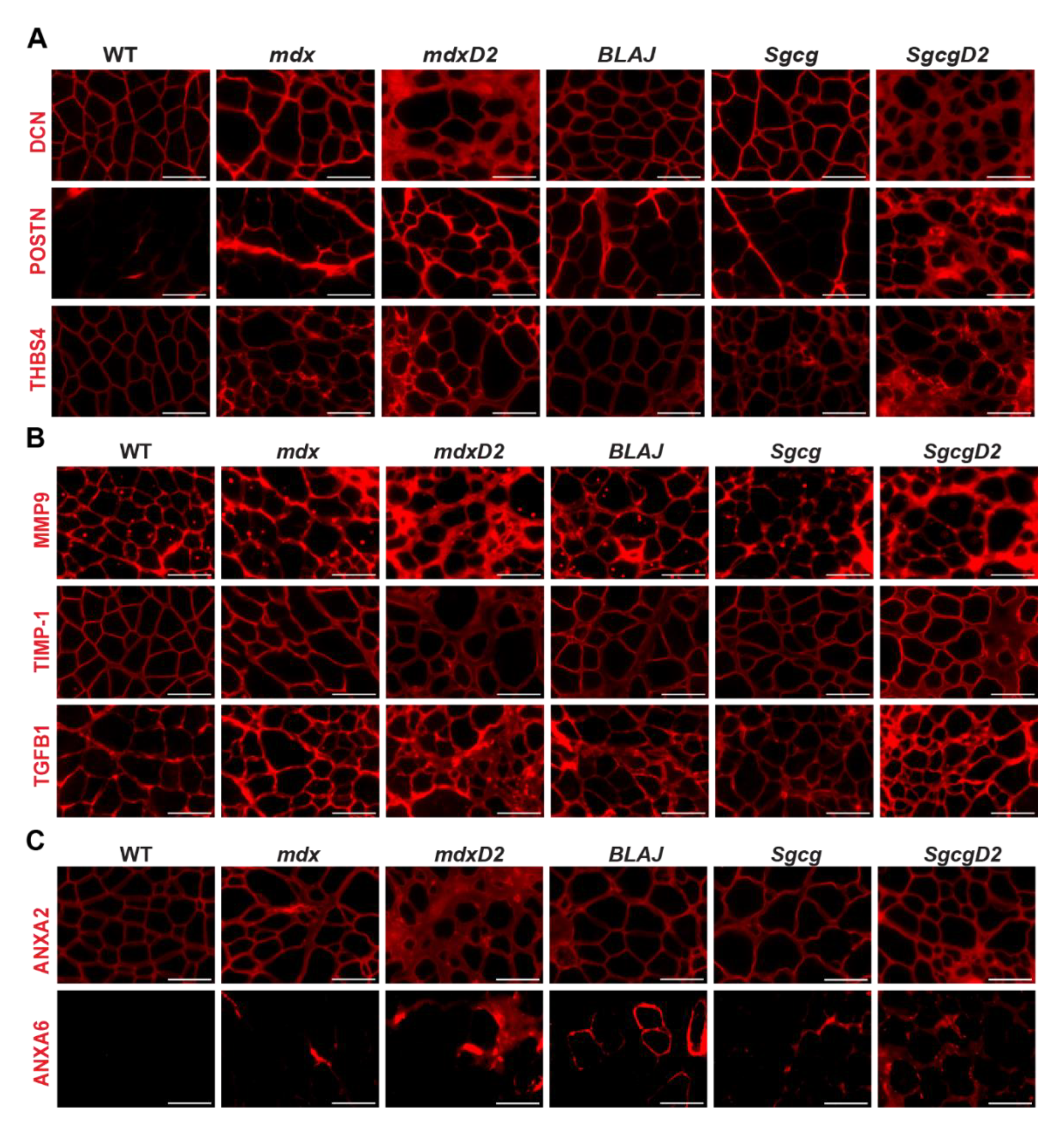
Representative IFM images from acellular myoscaffolds generated from WT, *mdx*, *mdxD2*, *BLAJ*, *Sgcg*, and *SgcgD2* mice. (**A**) Core Matrisome Proteins: DCN, POSTN, and THBS4. (**B**) Matrisome-Associated Proteins: MMP9, TIMP1, TGF- β1, ANXA2, and ANXA6.

**Supplemental Figure 5.**
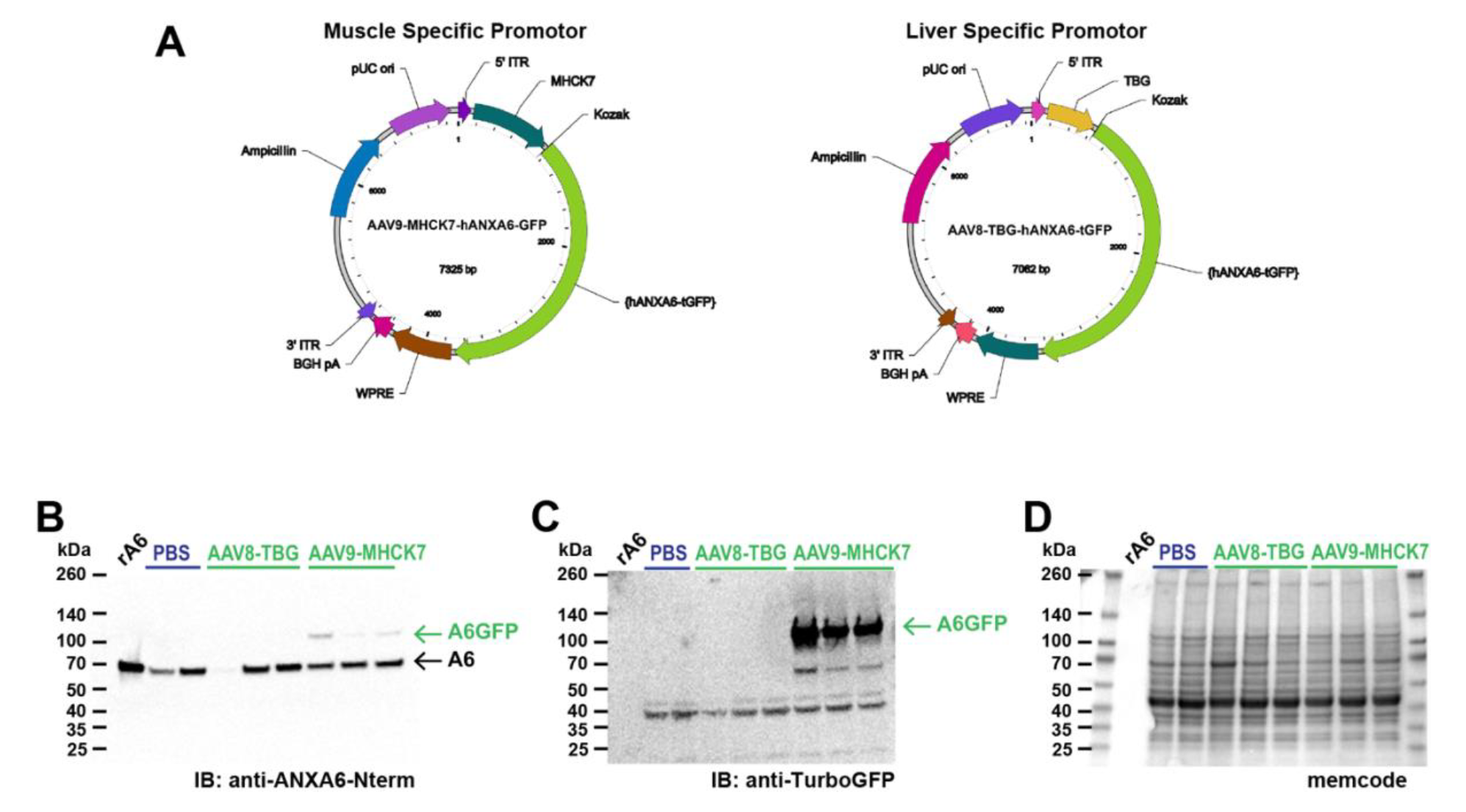
Adeno-asssociated expression of annexin A6-GFP within muscle. (**A**) Vector map of AAV9-MHCK7-hANXA6-GFP. This plasmid expresses GFP-tagged human annexin A6 driven by MHCK7, a striated muscle specific promoter, packaged in adeno-associated virus serotype 9 (AAV9). Vector map of AAV8-TBG-hANXA6-GFP. This plasmid expresses human annexin A6-GFP driven by TBG a liver specific promoter packaged in adeno-associated virus serotype 8 (AAV8). (**B-D**) Immunoblot of muscle lysates from PBS, AAV8- TBG, and AAV9-MHCK7 treated mice. Annexin A6-GFP (A6GFP) was present only in AAV9- MHCK7 muscle lysates evidence by the 100kDa anti-ANXA6 bands present in (**B**) and the anti-GFP bands present in (**C**). (**D**) Memcode illustrates equal loading across lanes.

**Supplemental Figure 6.**
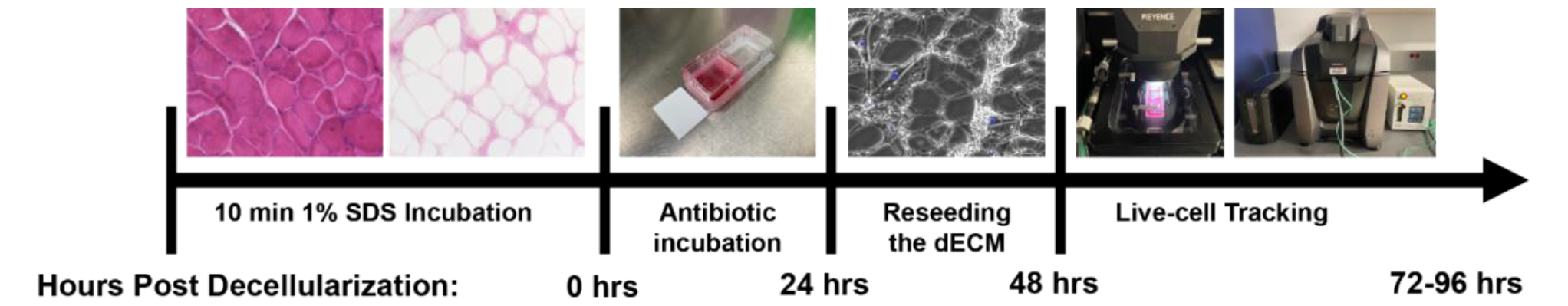
Diagram of the timeline for reseeding the acellular myoscaffold.

